# Characterization of cannabinoid plasma concentration, maternal health, and cytokine levels in a rat model of prenatal *Cannabis* smoke exposure

**DOI:** 10.1101/2023.06.16.545309

**Authors:** Tallan Black, Sarah L. Baccetto, Ilne L. Barnard, Emma Finch, Dan L. McElroy, Faith V. L. Austin-Scott, Quentin Greba, Deborah Michel, Ayat Zagzoog, John G. Howland, Robert B. Laprairie

## Abstract

*Cannabis sativa* has gained popularity as a “natural substance”, leading many to falsely assume that it is not harmful. This assumption has been documented amongst pregnant mothers, many of whom consider *Cannabis* use during pregnancy as benign. The purpose of this study was to validate a *Cannabis* smoke exposure model in pregnant rats by determining the plasma levels of cannabinoids and associated metabolites in the dams after exposure to either *Cannabis* smoke or injected cannabinoids. Maternal and fetal cytokine and chemokine profiles were also assessed after exposure. Pregnant Sprague-Dawley rats were treated daily from gestational day 6 – 20 with either room air, *i.p.* vehicle, inhaled high-Δ^9^-tetrahydrocannabinol (THC) (17.98% THC, 0.1% cannabidiol [CBD]) smoke, inhaled high-CBD (0.1% THC, 12.83% CBD) smoke, 3 mg/kg *i.p.* THC, or 10 mg/kg *i.p.* CBD. Our data reveal that THC and CBD, but not their metabolites, accumulate in maternal plasma after repeated exposures. Injection of THC or CBD was associated with fewer offspring and increased uterine reabsorption events. For cytokines and chemokines, injection of THC or CBD up-regulated several pro-inflammatory cytokines compared to control or high-THC smoke or high-CBD smoke in placental and fetal brain tissue, whereas smoke exposure was generally associated with reduced cytokine and chemokine concentrations in placental and fetal brain tissue compared to controls. These results support existing, but limited, knowledge on how different routes of administration contribute to inconsistent manifestations of cannabinoid-mediated effects on pregnancy. Smoked *Cannabis* is still the most common means of human consumption, and more preclinical investigation is needed to determine the effects of smoke inhalation on developmental and behavioural trajectories.

## Introduction

General trends of *Cannabis* legalization/decriminalization have changed social narratives surrounding use and risk perception in Canada and around the world. As social perception has changed, rates of use have increased for all ages by 7.3% since legalization, where smoking still accounts for 79% of *Cannabis* consumption in Canada [1]. Importantly, *Cannabis* has gained popularity as a “natural substance,” with the false assumption that it is therefore safe. Emerging human data indicate that children exposed to *Cannabis in utero* are at a higher risk of being born pre-term, underweight, and of developing persistent behavioural psychopathologies and cognitive deficits across their lifetime [2–8]. Although long-term outcomes are still not understood, these emerging data indicate the need for more preclinical research to establish the risks and investigate the molecular and physiological consequences of *in utero Cannabis* smoke exposure.

Over 120 unique phytocannabinoids are produced by the *Cannabis sativa* plant, with the acid forms of Δ^9^-tetrahydrocannabinol (THC) and cannabidiol (CBD) present at the highest relative concentrations [9]. These compounds, along with the lesser-known phytocannabinoids, terpenes, flavonoids, and alkaloids have a wide array of pharmacological activities via our endogenous cannabinoid system (eCS) and other biological systems [10]. The eCS is comprised of the G protein-coupled receptors (GPCR) cannabinoid 1 receptor (CB1R) and the cannabinoid 2 receptor (CB2R), their endogenous ligands [2-arachidonoylglycerol (2-AG) and anandamide (AEA)], and the enzymes that synthesize [diacylglycerolipase (DAGL) and N-arachidonoyl phosphatidyl ethanolamine-specific phospholipase D (NAPE-PLD)] and degrade [monoacylglycerol lipase (MAGL) and fatty acid amide hydrolase (FAAH)] those endogenous ligands, respectively [11].

THC, the main intoxicating constituent of *Cannabis*, accounts for the distinct “high” associated with *Cannabis* use via CB1R partial agonism [12]. CBD, by comparison, acts *in vitro* as a possible negative allosteric modulator at CB1R, partial agonist at CB2R, positive allosteric modulator of µ-opioid receptor, partial agonist of serotonin 1a receptors, and a modulator of the signaling of several other GPCRs and non-GPCR targets[13–15]. Evidence suggests that most of CBD’s *in vivo* activity occurs at non-cannabinoid receptor targets, such as the serotonin 1a receptor and the transient receptor potential vanilloid 1 (TRPV1) [16], further contributing to the pharmacological complexity of *Cannabis*.

Pregnancy is in part orchestrated by the intricate communication and coordination of maternal and fetal immune cells by cytokines and chemokines to establish and maintain the maternal-fetal interface [17]. Cytokines and chemokines play an important role in embryogenesis through the mediation of cell survival, proliferation, differentiation, and promotion of cell adhesion to specific sites to coordinate invasion of fetal trophoblasts into the maternal decidua [18–21]. Dysregulation in the establishment of a healthy placenta can result in maternal and fetal pathologies such as pre-eclampsia, spontaneous pre-term birth, intra-uterine growth restrictions, and resorption [22–26].

The eCS plays a formative role in the coordination of pregnancy and fetal neurodevelopment. Decreasing levels of intra-uterine AEA and concomitant signaling via CB1R and CB2R may dictate successful implantation [27], and changes in receptor expression and AEA throughout reproductive tissue and plasma has been associated with ectopic pregnancy, miscarriage and pre-eclampsia [28]. CB1R is also thought to coordinate directional migration and elongation of axons [29,30], and the maturation of astrocytes and oligodendrocytes [31–33]. THC and its constituents cross the blood-placenta barrier to accumulate in the developing fetus [34–37]. THC administration *in utero* interferes with endogenous cannabinoid orchestration of neuronal growth cones, cytoskeletal architecture, and the development of novel synapses [38], as well as in increased incidents of fetal growth restriction [39]. Existing preclinical studies of the downstream effects of gestational cannabinoid exposure have been performed using intravenous (*i.v.*), oral, intraperitoneal (*i.p.*), and subcutaneous (*s.c.*) administration methods of THC (0.5 mg/kg - 30 mg/kg) [36,39–45] and high-potency synthetic CB1R agonists (*e.g.* WIN-55,212-2) [46–54], with some recent studies examining vapour THC inhalation (100 ng/mL) and high-THC *Cannabis* smoke [37,55–57].

Plasma cannabinoid pharmacokinetics have been studied in healthy adult humans for multiple forms of consumption[58–61]; however, these metrics are poorly characterized in pregnancy [36]. Reported peak serum concentrations of THC after smoking a *Cannabis* joint achieve *t*_max_ approximately 12 min after inhalation and vary widely (C_max_ 60-200 ng/mL), due to inter-individual variability, variation in THC concentrations and frequency of use, among other factors [58–61]. Our ability to make conclusions based on human studies is therefore limited. In human studies, *Cannabis* consumption by expectant mothers is often measured by the number of joints smoked per day [36]. This method of measuring *Cannabis* consumption in humans makes comparing cannabinoid doses and exposures between clinical assessments and pre-clinical smoke exposure models challenging. Whereas prior preclinical studies have explored injection, inhalation, and oral administration of isolated THC or CBD [34–37,55,56,62–65], administering THC and/or CBD alone fails to take into consideration the potential effects of other phytocannabinoids, terpenes, and flavonoids of the *Cannabis* plant as well as the smoke of a combusted product [10,66,67]. Our group has previously worked to establish rat models of *Cannabis* smoke exposure in the contexts of absence epilepsy [82], cognition[68], and prenatal exposure during pregnancy on offspring health and behaviour [57]. In the present study, we build on this previous work to directly compare the effects of repeated inhalation of smoke from *Cannabis* products high in THC or CBD to repeated injections of THC or CBD in pregnant rats. Specifically, the pharmacokinetics of THC, CBD, and their metabolites; as well as the effects of *Cannabis* exposure on an array of 27 cytokines and chemokines was measured in pregnant Sprague-Dawley rats and their fetuses treated with *Cannabis* or cannabinoids daily from gestational day (GD) 6 – 20. The aim of our studies was a validation of a whole plant combustion method in gestation to enhance translation and address unanswered questions regarding the effects of *Cannabis* exposure during gestation.

## Results

### Pharmacokinetic evaluation and comparison of maternal plasma phytocannabinoid levels during gestation: accumulation of cannabinoids over time

Pregnant dams were treated daily between GD6 – 20 with either vehicle (*i.p.*), room air, 3 mg/kg *i.p.* THC, 10 mg/kg *i.p.* CBD, Skywalker Kush *Cannabis* flower smoke (high-THC smoke, 18% THC:0.1% CBD) 300 mg inhaled, or Treasure Island *Cannabis* flower smoke (high-CBD smoke, 0.1% THC:13% CBD) 300 mg inhaled (Fig. 1). Maternal plasma samples were collected from separate groups of rats 30 min after the first (GD6) and last (GD20) treatment to evaluate maternal phytocannabinoid levels (Fig. 1-3). For specific details on *Cannabis* combustion protocol refer to “*Smoke exposure”* in the Methods section below.

**Figure 1.**
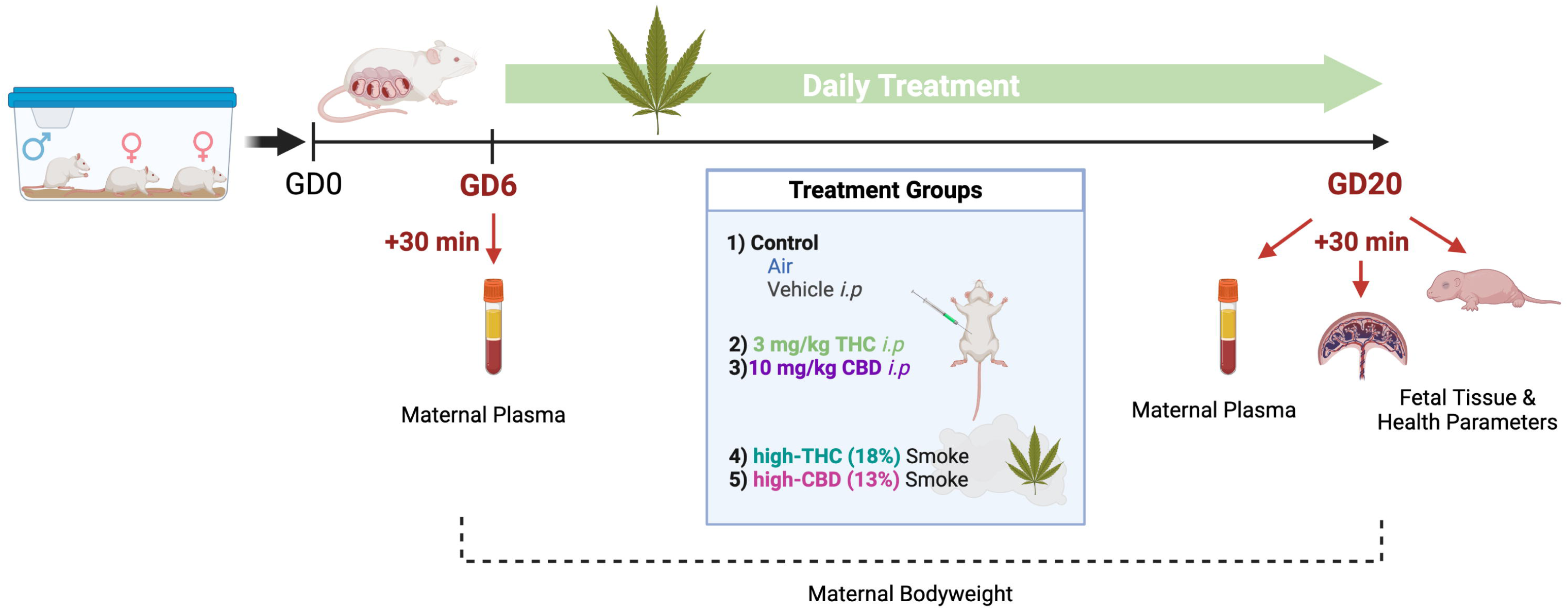
Experimental timeline for gestational *Cannabis* exposure. Rats were bred and treated once daily between gestational day (GD) 6 and 20 for 15 consecutive days of treatment. Maternal and fetal tissues were harvested 30 min following the first (GD6) and last (GD20) treatment. THC, CBD, and their metabolites were quantified in maternal plasma using high performance liquid chromatography tandem mass spectroscopy (HPLC-MS/MS). Protein samples from placenta and whole fetal brain were prepared for cytokine and chemokine quantification. Created with Biorender.com.

THC, CBD, and their metabolite levels were compared between GD6 and GD20 within treatments to assess whether levels changed across the gestational period (Fig 2). For 3 mg/kg *i.p.* THC exposed dams, mean plasma levels of THC, 11-OH-THC, and 11-COOH-THC were above the lower limit of quantification (LLOQ) on GD6 and GD20. A 3×2 ANOVA (Analyte by Day) demonstrated a significant main effect of Analyte (F_(2,30)_=5.70, p=0.008), but not Day (F_(1,30)_=3.41, p=0.08), and no interaction (F_(2, 30)_=2.93, p=0.07). Holm-Šídák’s multiple comparisons test revealed a significant difference between plasma levels of THC on GD6 vs. GD20 in *i.p.* THC exposed dams, with higher levels on GD20 (p=0.015) (Fig. 2A).

**Figure 2.**
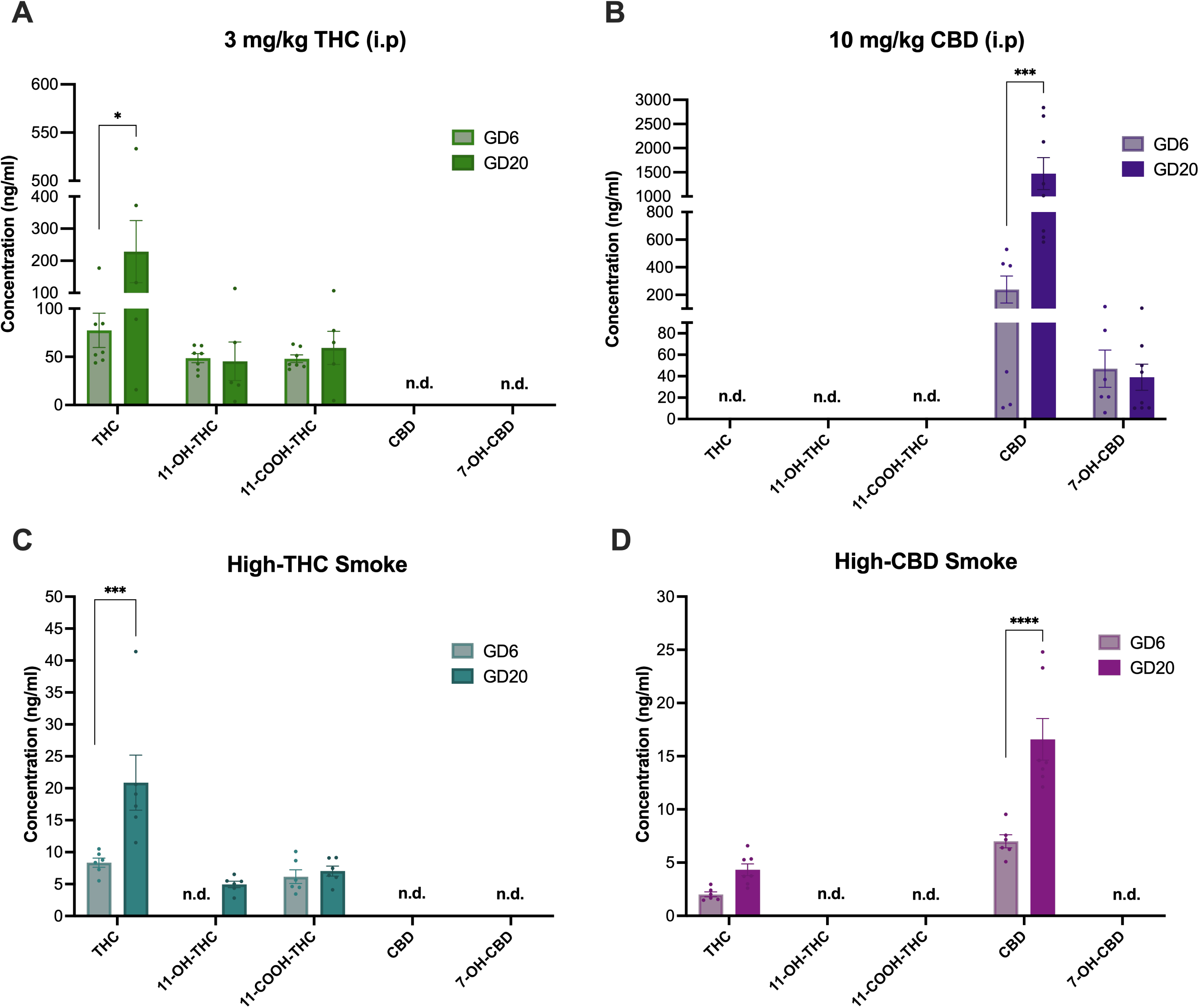
Comparison of THC, CBD, and metabolite levels in maternal plasma according to treatment between first exposure on GD6 and last exposure on GD20. Plasma levels of phytocannabinoids were quantified on GD6 and GD20 30 min after cannabinoid exposure in the following treatment groups: **(A)** 3 mg/kg/d THC *i.p.*, **(B)** 10 mg/kg/d CBD *i.p.,* **(C)** High-THC smoke, and **(D)** High-CBD smoke. Note the different scaling of the y-axes of the panels. Data are mean ± S.E.M. *n=*5-8 dams/per treatment. Values falling below LLOQ were deemed not detectable (n.d.).

**Figure 3.**
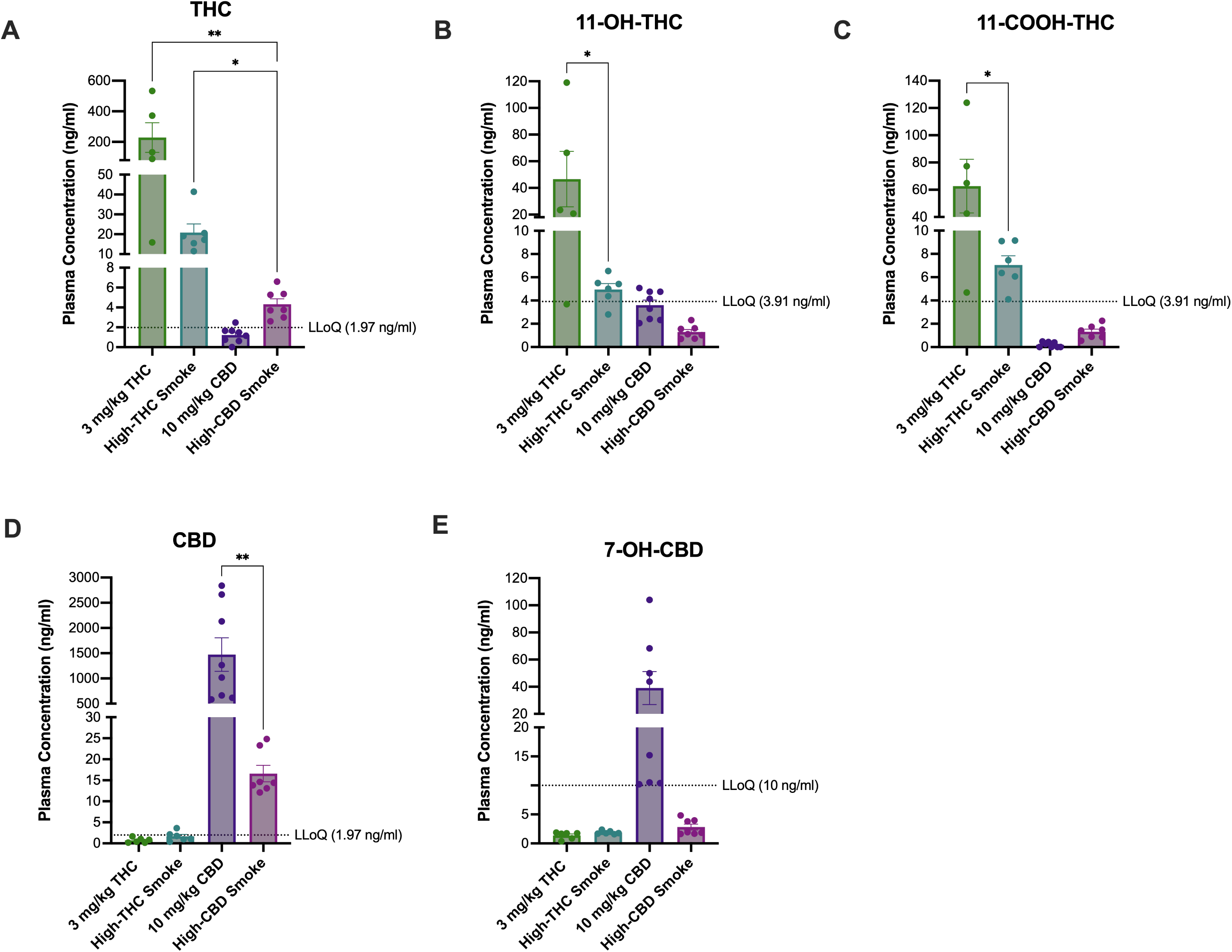
Comparison of THC, CBD, and metabolite levels among treatments in maternal plasma on GD20 after repeated exposure to either 3 mg/kg/d THC *i.p.*, high THC smoke, 10 mg/kg/d CBD *i.p.*, or high CBD smoke. Mean maternal plasma concentrations of **(A)** THC, **(B)** 11-OH-THC, **(C)** 11-COOH-THC, **(D)** CBD, and **(E)** 7-OH-CBD. All samples were collected on GD20, 30 minutes following final treatment. Note the different scaling of the y-axes of the panels. Data are mean ± S.E.M. *n=*5-8 dams/treatment; *p<0.05, ** p<0.01 as determined by Kruskal-Wallis (KW) one-way ANOVA or Kolmogorov-Smirnov (KS) test.

For 10 mg/kg *i.p.* CBD exposed dams, mean plasma levels of CBD and 7-OH-CBD were above the LLOQ. A 2×2 ANOVA (Analyte by Day), demonstrated significant main effects of Analyte (F_(1,24)_=16.83, p=0.0004) and Day (F_(1,24)_=9.57, p=0.005), as well as a significant interaction (F_(1,24)_=9.82, p=0.0045). Holm-Sidak’s multiple comparisons test revealed a significant difference between plasma levels of CBD on GD6 vs. GD20 in *i.p*. CBD exposed dams, with higher levels on GD20 (p=0.0004) (Fig. 2B).

For high-THC smoke exposed dams, mean plasma levels of THC and 11-COOH-THC were above the LLOQ on GD6 and GDG20; however, 11-OH-THC was only above the LLOQ on GD20. A 2×2 ANOVA (Analyte by Day) demonstrated significant main effects of Analyte (F_(1,20)_=12.36, p<0.0022) and Day (F_(1,20)_=8.64, p=0.0081), as well as a significant interaction (F_(1,20)_=6.52, p=0.019). Holm-Šídák’s multiple comparisons test revealed a significant difference between plasma levels of THC on GD6 vs. GD20 in high-THC smoke exposed dams, with higher levels on GD20 (p=0.0018) (Fig. 2C).

For high-CBD smoke-exposed dams, mean plasma levels of THC and CBD were above the LLOQ. The 2×2 ANOVA (Analyte by Day) demonstrated significant main effects of Analyte (F_(1,22)_=56.59, p<0.0001) and Day (F_(1,22)_=26.91, p<0.0001), as well as a significant interaction (F_(1,22)_=10.07, p=0.004). Holm-Šídák’s multiple comparisons test revealed a significant difference between plasma levels of CBD only on GD6 vs. GD20 in high-CBD exposed dams, with higher levels on GD20 (p<0.0001) (Fig 2D).

From these data, we conclude that circulating levels of THC or CBD, but not their respective metabolites, increased by approximately 3-fold between GD6 to GD20 in all treatments, regardless of administration method.

### Pharmacokinetic evaluation and comparison of maternal plasma phytocannabinoid levels during gestation: route of exposure influences observed concentrations

Mean GD6 and GD20 plasma levels of THC, CBD, and their metabolites were compared among treatments. As these analyses revealed similar differences among the treatments at each timepoint, we depicted the data from GD20 in Fig. 3 and the data from GD6 in Supp. Fig 1. For GD20 samples, THC levels were above the LLOQ for *i.p* THC, high-THC smoke, and high-CBD smoke. The Kruskal-Wallis (KW) test revealed a significant difference among treatments (H(3)=13.70, p<0.00010), where the *post-hoc* showed that levels of THC were significantly higher following *i.p* THC and high-THC smoke than high-CBD smoke (Fig. 3A). It is noteworthy that mean levels of THC in the *i.p.* THC-exposed dams are approximately 10-fold higher than those of the high-THC smoke group; however, due to unequal variances between treatments, this difference did not emerge as significant in the analysis. Mean plasma levels of 11-OH-THC were above the LLOQ for the *i.p* THC and high-THC smoke groups only (Fig. 3B). Levels of 11-OH-THC in the *i.p* THC treatment were significantly higher than levels in the high-THC smoke group (KS: p=0.047). Mean plasma levels of 11-COOH-THC were greater than the LLOQ in the *i.p.* THC and high-THC smoke treatments only (Fig. 3C). Plasma levels of 11-COOH-THC were higher in the *i.p.* THC group compared to high-THC smoke (KS: p=0.047) (Fig. 3C).

Plasma levels of CBD were greater than the LLOQ for the *i.p.* CBD and high-CBD smoke groups only and were significantly higher with *i.p.* CBD compared to the high-CBD smoke-exposed group (KS: p=0.00030) (Fig. 3D). 7-OH-CBD was only detectable above the LLOQ in rats of the *i.p.* CBD group (Fig. 3E).

From these data, we conclude that exposure of pregnant dams to high-THC smoke was associated with quantifiable levels of THC, 11-OH-THC, and 11-COOH-THC that were approximately 10% of the respective levels of THC, 11-OH-THC, and 11-COOH-THC in pregnant dams treated with 3 mg/kg/d THC *i.p*. (Fig. 3A-C). Similarly, treatment with high-CBD smoke was associated with quantifiable levels of CBD on GD20. Levels of CBD and 7-OH-CBD in high-CBD smoke-exposed dams were 1% and 10% of the respective plasma levels observed in pregnant dams treated with 10 mg/kg/d CBD *i.p.* (Fig. 3D, E). Therefore, the smoke exposure paradigm used in these studies yields quantifiable levels of plasma cannabinoids in pregnant dams at levels lower than those observed with conventionally used rodent injection models when collected 30 min after the final treatment.

### Fetal resorptions and litter size – but not other maternal health outcomes – were affected by phytocannabinoid injection

Maternal weight gain was measured every other day beginning on GD0. Uterine resorptions, litter sizes, fetal weight, fetal brain to body weight (BW), and fetal bodyweight to placenta ratios were collected on GD20. For maternal weight gain (Fig. 4A), there was a significant main effect of Time (F_(2.11, 57.57)_= 527.7, p<0.00010) but no main effect of Treatment (F_(4,28)_=0.34, p=0.85) or Time by Treatment interaction (F_(28, 191)_=0.93, p=0.57). There was a significant difference in fetal-placenta resorptions between treatments (one-way Kruskal-Wallis test: H(5)=15.32, p<0.0041), where the *post-hoc* demonstrated that *i.p.* CBD treatment caused significantly more resorptions than control and high-CBD smoke exposure, but not *i.p.* THC (Fig. 4B). Litter sizes were smaller for dams in the *i.p.* THC group compared to control animals (one-way ANOVA followed by Holm-Šídák *post-hoc* analysis; F_(4,28)_=4.50, p=0.0062) (Fig. 4C). There was no difference in fetal BW between treatments (F_(4, 29)_=2.14), p=0.10) (Fig. 4D). No differences were observed between treatment groups in mean fetal BW (Fig. 4D), brain to BW ratio (F_(4, 27)_=1.07, p=0.39) (Fig. 4E), or BW to placenta ratio (F_(4, 29)_=1.37, p=0.27) (Fig. 4E). No differences in the ratios of male to female offspring were observed among the treatments (F_(4, 25)_=1.56, p=0.22) (Fig. 4G).

**Figure 4.**
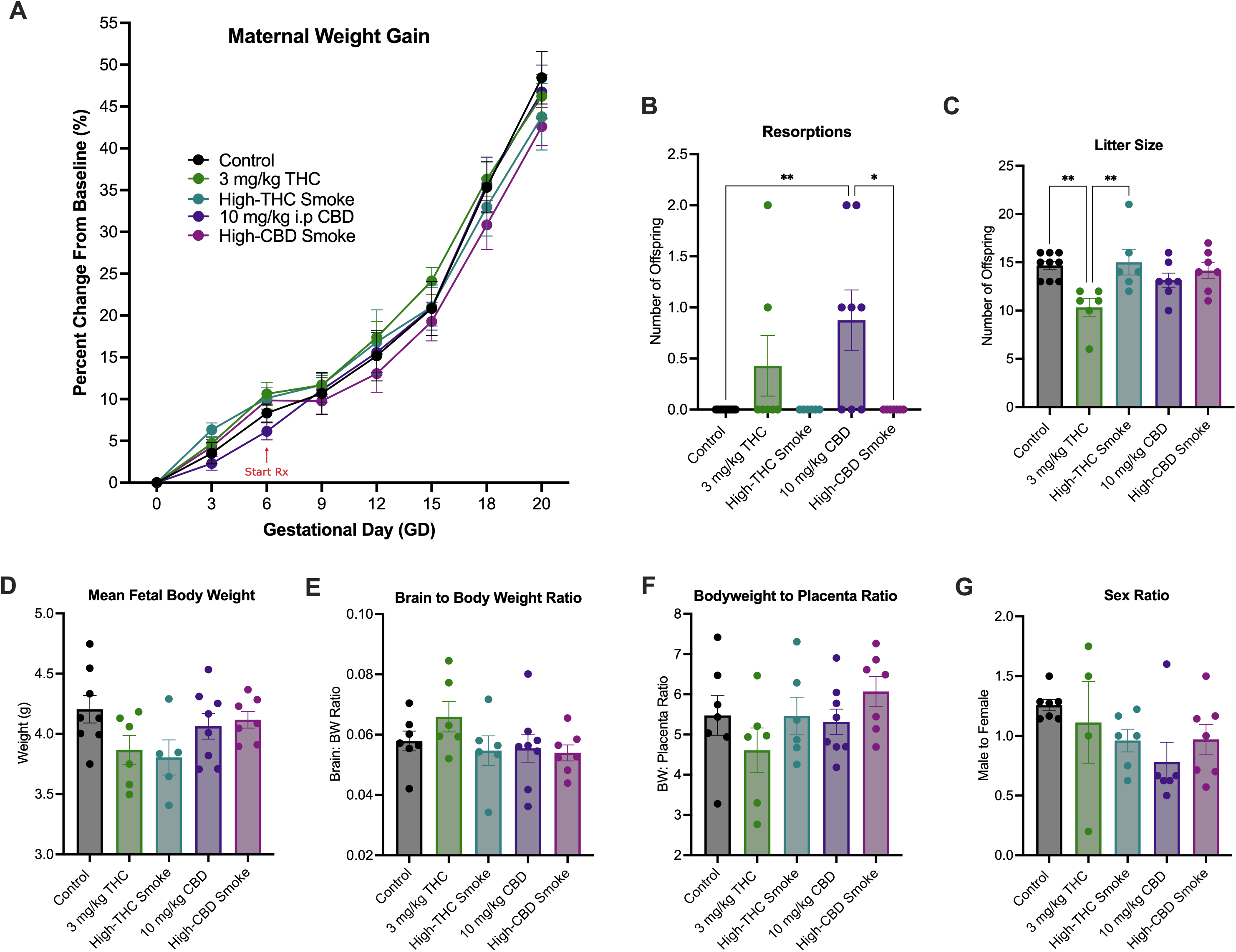
Markers of maternal and litter health during gestation. **(A)** Rates of maternal weight gain, **(B)** uterine resorption discovered, **(C)** litter size **(D)**, mean fetal body weight per litter, (**E)** mean fetal brain to body weight ratio per litter, and **(F)** mean fetal bodyweight to placenta ratio per litter, and **(G)** Ratio of male to female offspring per litter. Data are mean ± S.E.M, *n=*4-8 dams/per treatment. Measurements in panels B-G were taken on GD20. *p<0.05, as determined by one-way ANOVA followed by Holm-Šídák *post-hoc* analysis.

### THC injection was unique among treatment groups increasing or maintaining cytokine and chemokine levels relative to controls whereas other treatments were associated with reduced cytokine and chemokine levels

To survey the inflammatory state of placental and fetal tissue after phytocannabinoid treatment, 27 cytokines and chemokines were quantified in placental and fetal brain tissue collected on GD20, 30 min after final treatment.

#### Placenta

Significant main effect differences in cytokine or chemokine concentrations were observed for 15 of 27 cytokines and chemokines {epidermal growth factor (EGF); eotaxin; granulocyte colony-stimulating factor (G-CSF); interferon γ (IFNγ); IFNγ-induced protein 10 (IP-10, CXCL10); interleukins (IL) IL-1α, IL-1β, IL-4, IL-6, IL-10, and IL-17A; lipopolysaccharide-Induced CXC chemokine (LIX) or (CXCL5); macrophage inflammatory protein [MIP-1α; C-C motif chemokine (CCL3)], regulated up activation, normal T cell expressed and presumably secreted (RANTES, CCL5); and tumor necrosis factor α (TNFα)} (Suppl. Table 1). *Post-hoc* testing detected significant differences in 12 of 27 cytokines and chemokines (Fig. 5; Suppl. Table 1). Differences were not detected between groups in placental tissue for the remaining 15 cytokines and chemokines (Suppl. Table 1). The following is a summary of statistically significant *post-hoc* differences observed for cytokine and chemokine concentrations for placental tissue (Fig. 5). Data have been organized as being broadly defined as pro-inflammatory (red shape), anti-inflammatory (blue shape), or both depending on context (encapsulated by red and blue) (Fig. 5) [69,70].

**Figure 5.**
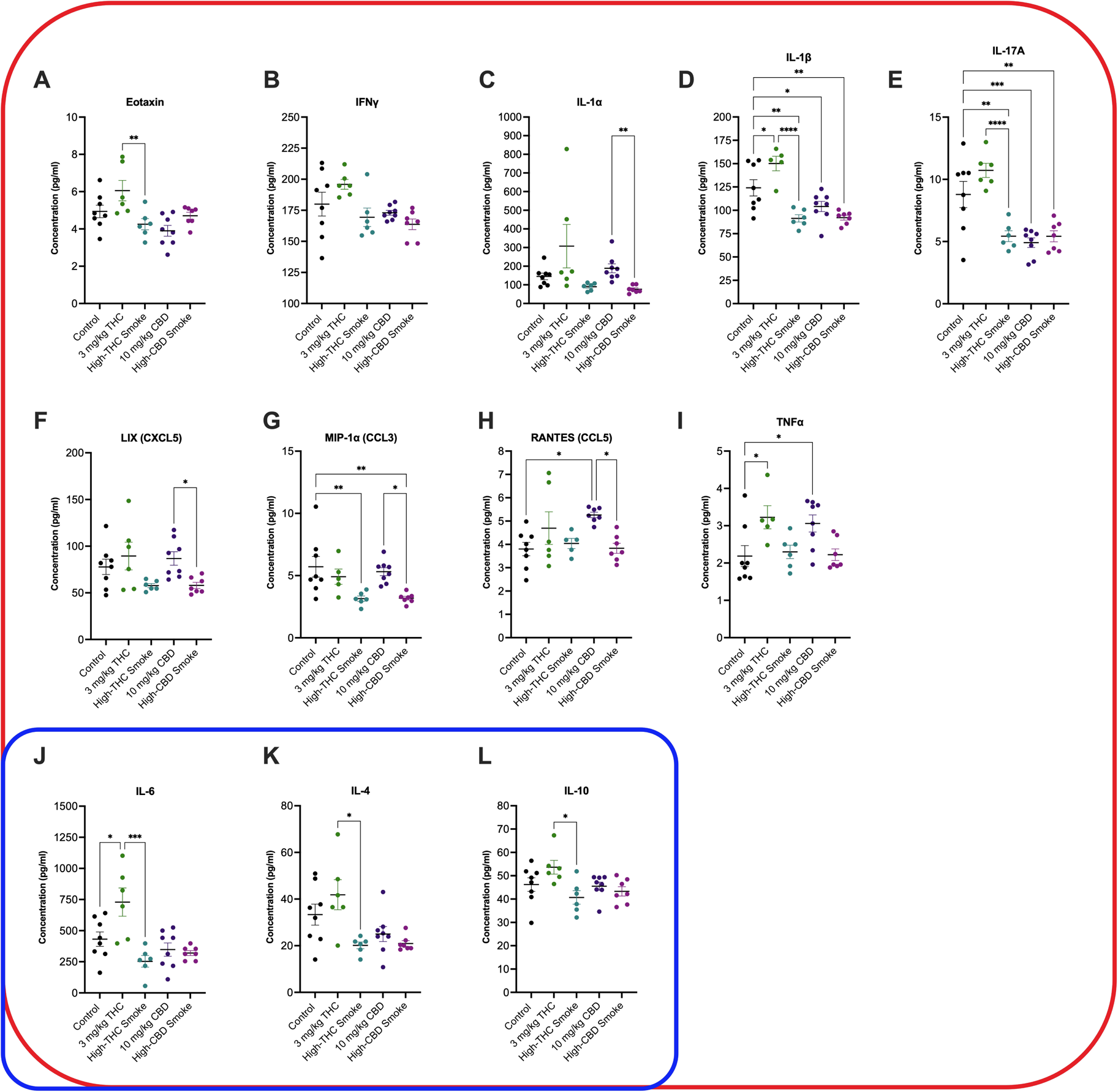
Placental cytokine and chemokine levels 30 min after final treatment on GD20. Cytokines and chemokines are organized by type: species within the red squircle are broadly considered pro-inflammatory, species within the blue squircle are broadly considered anti-inflammatory. Because the role of IL-4, IL-6 and IL-10 can be pro- or anti-inflammatory depending on context, these species are within both squircle. Data are displayed as mean ± S.E.M, *n*=6-8 dams/group. An average of two replicates was used when possible. Note the y-axis varies in each panel depending on the cytokine measured. *p<0.05, **p<0.01, ***p<0.001, ****p<0.0001 as indicated between groups.

Compared to control, the pro-inflammatory cytokines IL-1β (p<0.05) and TNFα (p<0.05) were elevated in placenta of dams receiving injected THC (Fig. 5D,I). Likewise, RANTES (p<0.05) and TNFα (p<0.05) were elevated in the placenta of dams receiving injected CBD (Fig. 5H,I). In contrast – and compared to control – the pro-inflammatory cytokines and chemokines IL-1β (p<0.01 for high-THC smoke and high-CBD smoke, p<0.05 for CBD injection) and IL-17A (p<0.01 for high-THC smoke and high-CBD smoke, p<0.001 for CBD injection), were decreased in the placenta of dams receiving high-THC smoke, high-CBD smoke, and injected CBD (Fig. 5D,E). Levels of the pro-inflammatory chemokine MIP-1α were also decreased in the placenta of dams receiving high-THC smoke or high-CBD smoke compared to control (p<0.01, Fig. 5G). Four differences were observed in levels of pro-inflammatory cytokines and chemokines when placenta from dams receiving injected THC were compared to other groups. Eotaxin (p<0.01), IFNγ (p<0.001), IL-1β (p<0.0001), and IL-17A (p<0.0001) levels were higher in the placenta of dams receiving THC injections compared to the placenta of dams receiving high-THC smoke (Fig. 5A,B,D,E). Similarly, IL-1α (p<0.01), LIX (p<0.05), MIP-1α (p<0.05), and RANTES (p<0.05) were all higher in the placenta of dams receiving CBD injections compared to the placenta of dams receiving high-CBD smoke (Fig. 5C,F-H).

IL-4, IL-6 and IL-10 are pro- and anti-inflammatory cytokines depending on how they interact with various cell types in pregnancy and other conditions [25]. In this study, IL-6 levels were elevated in the placenta of dams receiving THC injections compared to control (p<0.01) or compared to dams receiving high-THC smoke (p<0.001) (Fig. 5J). IL-4 (p<0.05) and IL-10 (p<0.05) levels were elevated in the placenta of dams receiving THC injections compared to dams receiving high-THC smoke (Fig. 5K & 5L).

From these data, three general trends emerge: (i) the majority of changes observed were for pro-inflammatory and not anti-inflammatory cytokines; (ii) when comparing to controls, injection of either THC or CBD was associated with the upregulation of four species (IL-1β, RANTES, TNFα, and IL-6), three of which are pro-inflammatory, whereas smoke exposure was generally associated with decreases in pro-inflammatory species; and (iii) upregulation of a subset of cytokines (eotaxin, IFNγ, IL-1β, IL-4, IL-6, IL-,10 IL-17A, and TNFα) was observed in placenta tissue of rats receiving THC injection relative to high-THC smoke and three species (LIX, MIP-1α, and RANTES) were similarly up-regulated in rats receiving CBD injection relative to high-CBD smoke. Together, these data highlight the widespread changes in inflammatory state that occurred in both injection and inhalation rodent models of prenatal cannabinoid exposure *and* illustrate that those injection and smoke exposure produced disparate outcomes in placental tissue.

#### Fetal brain

Significant main effect differences in cytokine or chemokine concentrations were observed for 16 of 27 cytokines and chemokines [EGF; IFNγ; IP-10 (CXCL10); ILs IL-1α, IL-1β, IL-2, IL-4, IL-6, IL-10, IL-17A, and IL-18; LIX (CXCL5); MIP-1α (CCL3); RANTES (CCL5); TNFα; and vascular endothelial growth factor (VEGF)] (Suppl. Table 2). *Post-hoc* testing detected significant differences in 15 of 27 cytokines and chemokines (Fig. 6; Suppl. Table 2). Differences were not detected between groups in fetal brain tissue for the remaining 12 cytokines and chemokines (Suppl. Fig. 2; Suppl. Table 2). The following is a summary of statistically significant *post-hoc* differences observed for cytokine and chemokine concentrations for fetal brain tissue (Fig. 6). Data have been organized as being broadly defined as pro-inflammatory (red squircle), anti-inflammatory (blue squircle), or both depending on context (encapsulated by red and blue) (Fig. 6) [69,70].

**Figure 6.**
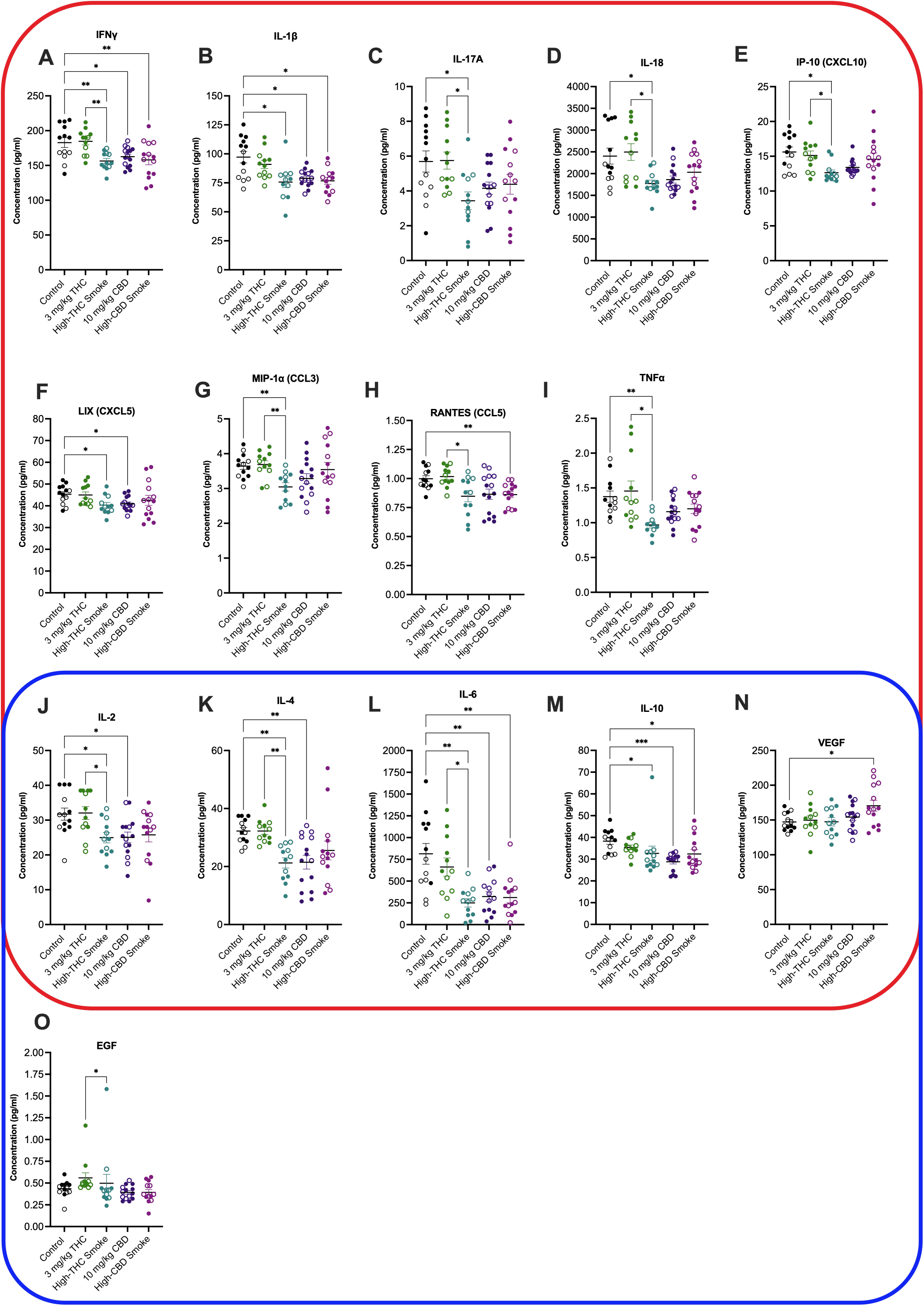
Fetal brain cytokine and chemokine concentrations 30 min after final treatment on GD20. Cytokines and chemokines are organized by type: species within the red squircle are broadly considered pro-inflammatory, species within the blue squircle are broadly considered anti-inflammatory. Because the role of IL-2, IL-4, IL-6, IL-10 and VEGF can be pro- or anti-inflammatory depending on context, these species are within both squircle. Data are displayed as mean ± S.E.M, *n*=12-15 samples/group. Closed circles are data points from males. Open circles are data points from females. An average of two replicates was used when possible. Note the y-axis varies in each panel depending on the cytokine measured. *p<0.05, **p<0.01 as indicated between groups.

Compared to control, the pro-inflammatory cytokines IFNγ (p<0.01 compared to high-THC or high-CBD smoke, p<0.05 compared to CBD injection) and IL-1β (p<0.01) were lower in fetal brains of dams receiving high-THC smoke, high-CBD smoke, and CBD injections (Fig. 6A,B). Levels of the pro-inflammatory cytokines IL-17A (p<0.05), IL-18 (p<0.05), IP-10 (p<0.05), LIX (p<0.05), and MIP-1α (p<0.01), TNFα (p< 0.01) were similarly lower in fetal brains of dams receiving high-THC smoke compared to control (Fig. 6C-E,G, I). LIX levels were also reduced in fetal brains from dams that received CBD injections (Fig. 6F); and RANTES levels were lower in fetal brains from dams that received high-CBD smoke compared to controls (p<0.01, Fig. 6H). Between treatment groups IFNγ (p<0.01), IL-1β (p<0.05), IL-17A (p<0.05), IL-18 (p<0.05), IP-10 (p<0.05), MIP-1α (p<0.01), RANTES (p<0.05) and TNFα (p< 0.05) were all elevated in fetal brains from THC injected grouped compared to high-THC smoke exposure groups (Fig. 6A-E, G-I).

IL-2, IL-4, IL-6, IL-10, and VEGF behave as pro- and anti-inflammatory cytokines depending on cellular context [70] . In this study, IL-2 (p<0.05) and IL-4 (p<0.01) levels were lower in fetal brain tissue of dams receiving high-THC smoke or CBD injections compared to control (Fig. 6J,K). Likewise, IL-6 (p<0.01) and IL-10 (p<0.05 for high-THC and high-CBD smoke, p<0.001 for CBD injections) were lower in fetal brain tissue of dams treated with high-THC smoke, high-CBD smoke, or CBD injections (Fig. 6L,M). VEGF levels were elevated in fetal brain tissues exposed to high-CBD smoke compared to controls (p<0.05, Fig. 6N). IL-2 (p<0.05), IL-4 (p<0.01), and IL-6 (p<0.05) levels were higher in the fetal brain tissues of dams treated with THC injections compared to high-THC exposure (Fig. 6J-N).

EGF is considered an anti-inflammatory chemokine [71]. EGF levels were elevated in the fetal brain tissues of dams that received THC injections compared to dams that received CBD injections (p<0.05, Fig. 6O).

From the fetal brain samples two patterns arise: (i) exposure to *Cannabis* smoke (high-THC *or* high-CBD) or injected CBD was associated with widespread downregulation of several cytokines and chemokines in fetal brains; and (ii) several analytes were notably elevated in fetal brain tissue from THC-injected groups relative to high-THC *Cannabis* smoke, but no *post-hoc* elevation was observed when THC-injected sample means were compared to control means. These data further demonstrate the widespread changes in inflammatory state that occur in both injection and inhalation rodent models of prenatal cannabinoid exposure *and* present several instances where changes observed in the placenta are similarly observed in the fetal brain.

## Discussion

In the present study, pregnant female rats were treated daily from GD6 until GD20 with either a control (*i.p.* vehicle injection or room air), 3 mg/kg/d *i.p.* THC, 300 mg/d high-THC *Cannabis* smoke (SW), 10 mg/kg/d *i.p.* CBD, or 300 mg/d high-CBD *Cannabis* smoke (TI). Results indicate that the duration of exposure (Fig. 2) and route of administration (Fig. 3) significantly affect the levels of cannabinoids and their metabolites in maternal plasma 30 min following administration. Analyses of maternal and fetal physiology over the course of pregnancy suggest relatively subtle effects of these treatment regimes, except for some evidence of reduced fetal viability following repeated injections of either THC or CBD (Fig. 4). Following repeated treatment of the dams, levels of cytokines and chemokines in the placenta (Fig. 5) and fetal brain (Fig. 6) were also altered in a cannabinoid- and route-dependent manner. Taken together, our results suggest that the route of administration of cannabinoids in developmental studies is an important factor in determining the specific effects obtained. Thus, care should be taken to use methods of administration to closely resemble those of humans to improve construct validity of preclinical models.

### Comparing plasma levels of THC and CBD following repeated injections or smoke administrations to pregnant rats

Analyses of plasma levels of cannabinoids revealed that plasma levels of THC and CBD, but not their metabolites, were significantly higher when sampled on GD20, after 15 administrations, compared to GD6, after one administration (Fig 2). To the best of our knowledge, this is the first-time plasma cannabinoid levels have been directly compared between acute and repeated administrations during pregnancy in rats. While the potential for accumulation during pregnancy is interesting, these results may be explained by several factors including - but not limited to – homeostatic changes to the metabolism of cannabinoids during repeated exposures or binding and depositing of these cannabinoids on plasma proteins and other tissue during repeated dosing. It is noteworthy that levels of THC and CBD following acute exposure on GD6 were lower than typically observed in male, but not female, rats. Barnard and colleagues [68] reported plasma levels of THC of 125 ng/ml (vs. approximately 77 ng/ml for GD6 here) and 15 ng/ml (vs. approximately 8 ng/ml for GD6 here) 30 min following acute treatment of male rats with the same THC injection and high-THC smoke protocols used in the present study. Plasma levels of CBD following high-CBD smoke exposure were approximately 15 ng/ml in the same study [68], which is also considerably higher than the roughly 5 ng/ml reported following the first administration of high-CBD smoke to the pregnant rats in this study. However, Baglot and colleagues [62] reported sex differences in the plasma concentrations of injected THC (2.5 mg/kg, i.p.) with females showing levels similar to those reported herein, and males showing higher levels similar to the Barnard [68] study [68]. In a comparison of THC (dronabinol) Volcano® inhalation treatment to *i.p.* injection in adult male rats, Manwell and colleagues [81] reported comparable plasma levels of THC at 20 min and at 40 min time points between 1 mg/pad THC and 2.5 mg/kg *i.p.* THC to males injected in Barnard [68]. Using a different protocol of vaporized THC exposure, Baglot and colleagues found higher plasma levels of THC in male and female rats 30 minutes after inhalation than reported in the present study [62]. Notably, Sandini and colleagues [57] detected behavioural changes in adult offspring after repeated prenatal exposure to high-THC smoke using like those used in the present study. In another inhalation model, Jenkins and colleagues [72] observed suppressed oscillatory power and coherence with acute THC flower vapour exposure in male rats; however, neither Jenkins et al., [72] nor Sandini et al., [57] provide pharmacokinetic support for their models.

To the best of our knowledge, there are no previous reports of CBD accumulation during pregnancy and the direct comparison of plasma levels of CBD following injected CBD or inhalation of high-CBD smoke in this study is novel. We found maternal plasma CBD levels at 30 min following 10 mg/kg/d *i.p.* were 239 ng/mL on GD6, increasing to 1,473 ng/mL after 15 daily injections on GD20 (Fig 3). In the high-CBD smoke group, levels of CBD were dramatically lower than in injected CBD dams, where on GD6 plasma levels were 7 ng/ml which increased to 17 ng/mL on GD20 at 30 min after administration. Our injected CBD GD6 plasma values more closely align with acute vapor administration in female rats at 100 ng/mL and 10 mg/kg *i.p.* [73] whereas our GD20 injected CBD plasma values resemble those from Ochiai and colleagues (2021), where a single dose of 10 mg/kg *i.v.* of CBD resulted in a *C*_max_ in maternal plasma of ∼1200 ng/mL and 900 ng/mL in whole fetal tissue [74]. Of note, Ochiai and colleagues (2021) reported the half-life of injected CBD (10 mg/kg; i.v.) as 5 hours in pregnant mice [74], which casts doubt on significant accumulation from repeated once daily injections. Thus, directly comparing plasma cannabinoid levels by sex and reproductive status will be an important consideration for future research. In addition, assessing plasma levels of cannabinoids *without* treatment on GD20 would confirm whether accumulation of the cannabinoids was occurring over repeated exposures.

Our experimental design also allowed for a direct comparison of plasma levels of cannabinoids following either smoke exposure or injections. Levels of THC and CBD, as well as some of their metabolites, were significantly higher 30 min following repeated injections than repeated smoke exposure on GD20. In general, we chose to compare injected doses of THC and CBD commonly used in rodent models [38,39,42,75–78] with the smoke inhalation protocol developed in our laboratory[57,68,79]. The inhalation protocol resembles other ‘hotbox’ methods [72,80,81], and has been shown to result in acute behavioural and oscillatory effects [68,79], as well as long-term changes in the offspring following exposure during pregnancy [57]. Despite previous acute and chronic behavioural disturbances associated with inhalation methods, this is the first pharmacokinetic evaluation comparing acute (GD6) and chronic (GD20) exposure in a prenatal model of whole cannabis flower smoke exposure. As reported previously for injected THC and smoke exposure [68], levels of THC were lower following inhalation than injections. Determining the peak and total exposure is difficult with one sampling point and the passive exposure methods we used with smoke administration; however, it is likely that our smoke exposure protocol results in lower levels of exposure when compared to some other inhalation methods [37]. In addition, different pharmacokinetics following these administration methods may dramatically affect the peak and total exposure of the pregnant woman and fetus to cannabinoids.

### Relatively subtle effects of repeated cannabinoid administration on maternal and fetal parameters

Injected cannabinoids and inhaled *Cannabis* did not significantly impact maternal weight gain, fetal weight, fetal brain to BW or BW to placenta ratios, or sex ratios in this study. Previously, we observed no effect of repeated high-THC smoke exposures on maternal weight gain, litter size, or litter weight following high-THC smoke exposure [57], although in that study an increase in the ratio of male:female offspring was significant. A limitation of this study is the relatively small sample sizes for the treatment groups that were carried to GD20, which may have limited the emergence of more nuance differences in markers of maternal and fetal health. However, injected CBD did increase fetal resorptions and injected THC decreased litter size. Natale and colleagues (2020) reported increased rates of uterine resorption in mice treated with 3 mg/kg/d *i.p.* THC, alongside indicators of intra-uterine growth restriction (IUGR) and placenta abnormalities [39]. Although CBD has not been previously investigated in these models *in vivo*, *in vitro* assays of trophoblasts show that CBD has a dose-dependent negative impact on cell cycle progression, viability, and migration; all crucial mechanisms in fetal trophoblast development [82]. Analysis of clinical maternal and childhood health outcomes has yielded different results due to confounding variables such as polysubstance use and inaccurate reporting of *Cannabis* use frequency. However, clinical retrospectives and pre-clinical investigation have been associated with adverse neonatal outcomes such as decreased birthweight and APGAR scores, preterm births, miscarriages, growth restrictions, and placenta abnormalities (reviewed in [83]), and new epidemiological data show geospatial associations in the US between congenital abnormalities and *Cannabis* availability, specifically CBD products [84]. It is therefore possible that injected THC and CBD produce similar restrictions to healthy litter growth resulting in the loss of pups. Because neither smoke treatment used in the current study produced any differences in measures of maternal and fetal health, we suggest that injected and inhaled *Cannabis* models differ in their effects on maternal and fetal health.

### Repeated cannabinoid treatments differentially alter cytokine levels in placenta and fetal brain tissue

In GD20 placental tissues from rats repeatedly exposed to either THC or CBD, we observed significant differences in levels of 13 of 27 cytokines and chemokines compared to control (Fig. 5). Injected THC treatment upregulated 4 inflammatory cytokines and chemokines, whereas injected CBD only elevated TNFα (also observed with injected THC) and RANTES (not observed for injected THC). Smoke exposure, regardless of the cannabinoid content, generally decreased cytokine and chemokine levels. Analyses of fetal brain tissue harvested from the same pregnancies as the placental tissue revealed significant alterations in levels of 14 of 27 cytokines and chemokines tested among the groups (Fig. 6). Although the effects of maternal *Cannabis* exposure on immune mechanisms in the mother and offspring are of current interest [85], limited *in vivo* data exist regarding the maternal and fetal immune response to repeated cannabinoid treatments, and potential sex dependant differences. Our results show intriguing changes 30 min following the final treatment on G20. Determining the time course and significance of these changes for placental and fetal health will require additional experimentation. IL-1β, IL-6, IP-10 (CXCL10), and TNFα are all pro-inflammatory cytokines and chemokines important for macrophage and natural killer (NK) cell activation, where the presence of NK cells correlate with fetal resorption, due to their damaging effects on trophoblast proliferation [23]. Elevations in IL-6, TNFα, and IP-10 (CXCL10), specifically amongst placenta and fetal tissue, have been associated with preterm births, intrauterine fetal death, and resorptions in addition to other downstream pathophysiological and behaviour changes [86–88].

## Conclusion

Our results show that injection of THC or CBD during gestation produces markedly different physiological effects when compared to smoke exposure with high-THC or high-CBD *Cannabis*. Smoke exposure did not cause any large-scale changes in markers of maternal or fetal health but did change the levels of immunomodulatory cytokines and chemokines in both placenta and fetal brain samples. Further in-depth behavioural analyses of offspring following exposure to *Cannabis* smoke compared to injected cannabinoids is required. An important limitation of our current model is that cannabinoids were administered to rats during the equivalent of the late first and second trimester of humans, and therefore our exposure model does not reflect potential use patterns and effects during critical neuro-developmental periods before conception, early in gestation, or the third trimester. Additionally, our model of smoke inhalation uses passive exposure, which standardizes the exposure that dams received. However, active self-administration models (*e.g.*, [56]), may better represent voluntary usage patterns and reduce exposure stress. Overall, these data highlight the fundamental differences in outcomes that route, dose, and formulation of *Cannabis* can have. These results should be considered in our interpretations of rodent model data for gestation and other applications of smoke inhalation and cannabinoid injection.

## Methods

### Animals

Virgin female (n=76) and male Sprague-Dawley rats (n=24; Charles River, Senneville, QC) arrived at our animal housing facility and were allowed to habituate for at least 1 week in ventilated cages in a climate-controlled vivarium under a 12h:12h light-dark cycle (lights on at 07:00). *Ad libitum* food (Purina rat chow) and water was provided throughout all experiments. Animals were then handled for 1 week before males and females (8-10 weeks of age) were paired at 17:00 and left overnight to breed. Evidence of sperm following vaginal swab at 09:00 the following morning was considered GD0. Dams were left undisturbed until GD6, when drug treatments began (Fig 1). Dams were randomized and assigned treatment with either high-THC *Cannabis* smoke (Skywalker Kush [SW]: 17.98% THC, 0.1% CBD, Aphria Inc., Lot# 5142072190); high-CBD *Cannabis* smoke (Treasure Island Kush [TI]: 12.83% CBD, 0.68% THC, Aphria Inc., Lot# 5142071556); or *i.p.* injections of 3 mg/kg/d THC (Cayman Chemical, Ann Arbor, MI Cat. No. 9003740), 10 mg/kg/d CBD (Cayman Chemical, Cat. No. 9003741), vehicle (*i.p.*) (1:1:18 ratio ethanol:kolliphor:phosphate buffered saline) or air exposure once daily between GD6 - 20 (Fig 1). Once pregnant, dams were singly housed in standard cages with adequate cage enrichment. All experiments were conducted according to standards set by Canadian Council on Animal Care and the University of Saskatchewan Research Ethics Board (Protocols 20210009, 20190067).

### Smoke exposure

Pregnant rats (n=5-8/group) were treated using a well-validated inhalation system (La Jolla Research Inc.) and method established in [57,68,79]. *Cannabis* was shredded and ground using a coffee grinder (10 sec). Each bowl (ceramic coil) was packed with 300 mg of the ground *Cannabis* flower. Airtight chambers (33 cm *x* 30.5 cm *x* 51 cm) equivalent to approximately 50 L each, could house a maximum of 2 rats, separated into plastic cages with metal grate roof used exclusively in the smoke chambers. Rats were habituated to the smoke chamber and pumps for 2 days for 20 min prior to smoke exposure. Before combustion, rats were placed in the chambers with pumps turned on for 5 mins. Air was pumped through the chambers at 10 L/min and exhausted into a fume hood. *Cannabis* was combusted over a period of 1 min and the resulting smoke continuously pumped through the system. Pumps were turned off for 1 min following the combustion protocol before the smoke was vented gradually over 13 min.

### Tissue collection

Maternal blood collection was performed via cardiac puncture. Rats were anesthetized with 5% isoflurane, and maternal blood, organs and fetal tissues were collection 30 min following the start of their last drug treatment. Fetal and maternal tissues were placed in 5 mL LoBind Eppendorf tubes (Cat. No. 0030122356), frozen in liquid nitrogen, and stored at -80°.

Cardiac blood was collected by syringe and immediately transferred to 4 mL BD Vacutainer tube (Cat. No. CABD367884). All blood samples were centrifuged for 10 min at 4° at 2,000 *x* g under 1 h from time of collection. Plasma was collected, aliquoted at 300 µL into 1.5 mL Lobind Eppendor tubes (Cat. No.0030108442), frozen and stored at -80° until needed.

### HPLC-MS/MS

The pharmacokinetic investigation of maternal plasma concentrations of THC, CBD and their associated metabolites ±11-OH-Δ^9-^ THC (11-OH-THC), ±11-nor-9-carboxy-Δ^9-^ THC (11-COOH-THC), 6α-OH-CBD and 7-OH-CBD was performed using high performance liquid chromatography tandem mass spectrometry (HPLC-MS/MS) [60] (Fig 2). Phytocannabinoids were quantified using Agilent 1260 binary pump LC system (agilent Technologies Canada, Mississauga, Ontario, Canada) coupled to an ABSciex 4000QTRAP (ABSciex, Concord, ON) mass spectrometer using a Eclipse Plus phenyl hexyl column (4.6 A 100mm, 5 µm column, Agilent) and 1290 Infinity II inline filter (0.3 µm). HPLC-MS/MS was run and peaks analyzed using Analyst (version 1.7).

CBD, CBD-D3, THC, THC-D3, 11-OH-THC, 11-COOH-THC working and internal standards were purchased from Cerilliant, where 6α-OH-CBD and 7-OH-CBD and 7-OH-CBD-D9 were purchased from Toronto Research Chemicals and diluted to 1 mg/mL and stored at -20 ° . The standard curve was established between 39.06 ng/mL and 5,000 ng/mL. The LLOQ for THC and CBD was 1.97 ng/mL. For 11-OH-THC, 11-COOH-THC, and 6α-OH-CBD, and 7-OH-CBD the LLOQ were 10 ng/mL. All phytocannabinoids and their metabolites shared a medium quality control (MQC) of 100 ng/mL and a high-quality control of 175 ng/mL. All 6α-OH-CBD was below our LLOQ. Standard curve and quality controls were prepared by mixing 10 µL of standard dilutions to 190 µL of blank plasma. Six hundred µL of prepared internal standard solution at 1.84 µg/mL was added to all standards and QCs, vortexed and centrifuged at 14,000 *x* g at 4° to precipitate proteins. Supernatant was collected, vacuum filtered through an Agilent Captiva EMR-Lipid 96-well plate (Cat. No. 5190-1001) and transferred to an amber autosampler vial for testing. To prepare maternal plasma, samples were thawed at room temperature, 600 µL of internal standard solution was added, samples were vortexed, centrifuged and filtered as above.

### Cytokine and chemokine array

Whole fetal brains were dissected, and a transverse dissection of the placenta was collected and homogenized (Rotor-Stator, OMNI International) over ice using RIPA buffer (Cat. No.89901 ThermoFisher) and 1:100 HALT Protease and Phosphatase Inhibitors (Cat#:78440 ThermoFisher, Burlington, ON) at a ratio of 750 µL RIPA to 150 mg of tissue. Samples were centrifuged at 12,000 *x* g for 10 min at 4°C and the supernatant was transferred to a new Lobind Ependorf tube (Cat. No. 0030108442), this step was repeated twice. Protein concentrations were quantified using the Pierce^TM^ BCA protein assay kit (Thermofisher). Four thousand µg/mL of sample was aliquoted, and samples were analyzed using the Rat Cytokine 27-Plex (Millipore Milliplex) by Eve Technologies (Calgary, AB).

### Statistical analyses

All data analyses for this study were conducted and graphed with Prism (GraphPad, v. 9.0). The pharmacokinetic data (Fig 2 and 3) were analyzed two separate ways. One injected THC dam was removed from analysis on GD20 due to low plasma THC levels. Plasma cannabinoid concentrations were expressed as a mean ± S.E.M. *n*=5-8 litters per group. Only samples that were above the LLOQs were analyzed. Data in Fig. 2 and 3, as well as Supp Fig.1, were analyzed with ANOVA or nonparametric tests as appropriate (see Results for specific details). Percentage maternal weight gain across gestation was calculated as: ((Final-Baseline)/Baseline*100)). Data were analyzed with a 2 level (7 x 5: Day x Treatment) mixed effects model (REML), where data are expressed as a mean ± S.E.M., *n*=6-7 dams per treatment. Other maternal and pup health data were assessed for normality (KS-test) and homoscedasticity (Brown-Forsyth test) and analyzed with one-way ANOVA with Greenhouse-Geisser correction or Kruskal-Wallis (KW) test (non-parametric) with maternal Treatment as a between-subjects factor. Planned comparisons between control to injected-THC, control to injected-CBD, injected-THC to high-THC smoke, injected-CBD to high-CBD smoke, and high-THC to high-CBD smoke were performed using Holm-Šídák’s (parametric) or Dunn’s (non-parametric) multiple comparisons tests. The multiplex data were assessed for normality (KS-test) and homoscedasticity (Brown-Forsyth test). When data met these assumptions, they were analyzed according to an ordinary one-way ANOVA with Greenhouse-Geisser correction. If data violated the KS-test, outliers were identified, and data were further analyzed using ordinary one-way ANOVA with Greenhouse-Geisser correction and KW-test if not normally distributed. If data were heteroscedastic, Brown-Forsythe ANOVA was used. Planned comparisons (as listed above) were performed using Holm-Šídák’s (parametric), Dunn’s (non-parametric), or Dunnett’s T3 tests (heteroscedastic) to compare mean concentrations of each treatment group.

## Acknowledgements

The authors acknowledge Maddie Stewart’s for input and assistance regarding interpretation of the multiplex experiment.

## Author Contributions

TB planned and executed the experiment, analyzed all data, wrote, and edited the manuscript with the technical and practical assistance of SLB, ILB, EF, DLM, FVLA-S, and QG for animal work. DM and AZ provided assistance and oversight for the pharmacokinetic experiments and data analysis. JGH and RBL designed and planned the experiments, supervised all highly qualified personnel, assisted with data analysis, and contributed to the writing and editing of the manuscript.

## Additional Information

Supplementary information is available for this manuscript online including all statistical analyses and all data not presented directly in the body of the manuscript.

