## Supplementary Information for "Characterization of cannabinoid plasma concentration, maternal health, and cytokine levels in a rat model of prenatal *Cannabis* smoke exposure"

Abbreviated title: Rat prenatal *Cannabis* smoke exposure

^1^College of Pharmacy and Nutrition, University of Saskatchewan, Saskatoon SK Canada

^2^Department of Anatomy, Physiology, and Pharmacology, College of Medicine, University of Saskatchewan, Saskatoon SK Canada

^3^Department of Pharmacology, College of Medicine, Dalhousie University, Halifax NS Canada

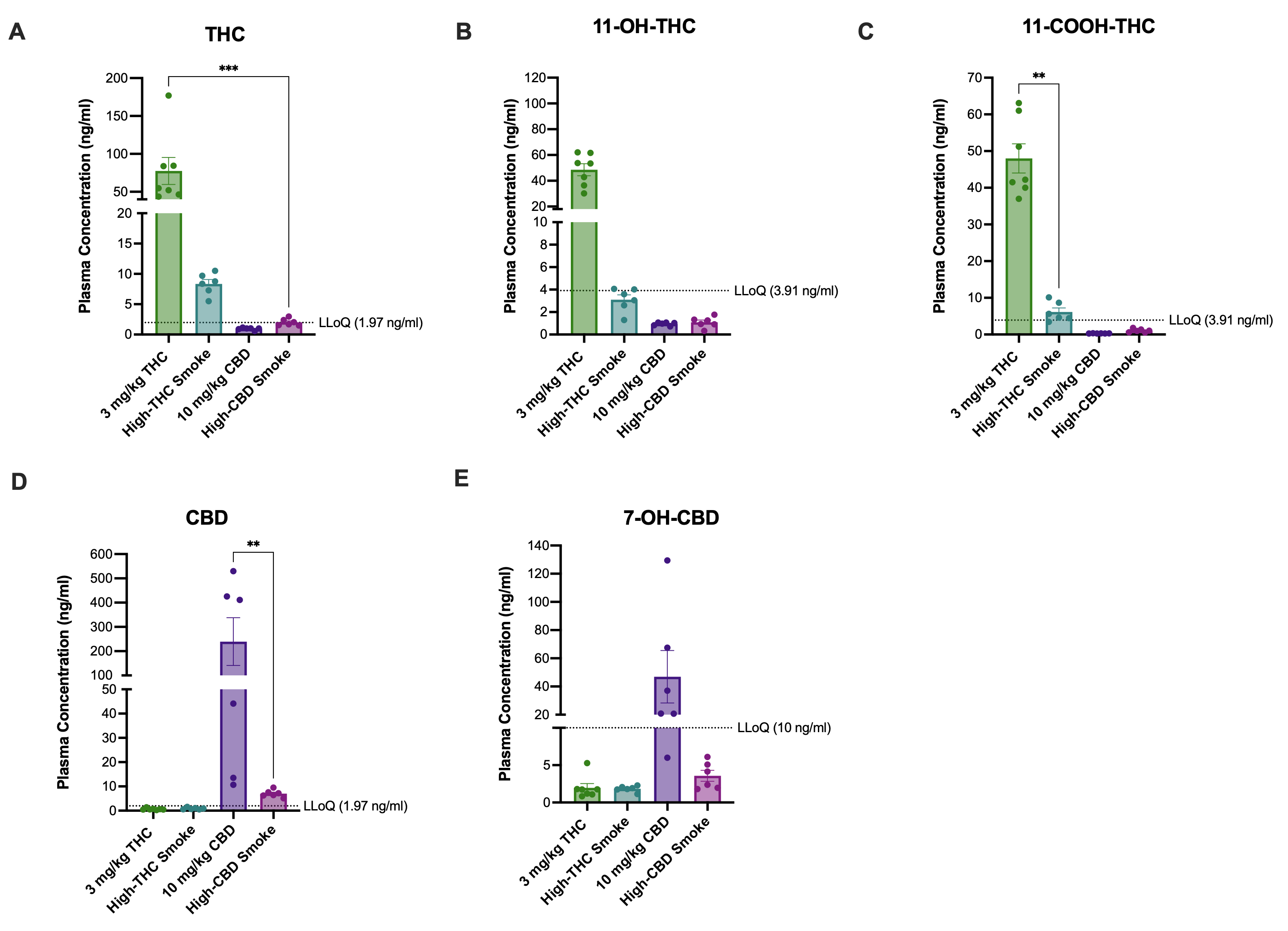

**Figure S1.** Comparison of THC, CBD, and metabolite levels among treatments in maternal plasma on GD6 after a single exposure to either 3 mg/kg/d THC *i.p.*, high THC smoke, 10 mg/kg/d CBD *i.p.*, or high CBD smoke. Mean maternal plasma concentrations of **(A)** THC, **(B)** 11-OH-THC, **(C)** 11-COOH-THC, **(D)** CBD, and **(E)** 7-OH-CBD. All samples were collected on GD6, 30 minutes following one treatment**.** Note the different scaling of the y-axes of the panels. Data are mean ± S.E.M. *n=*6-8 dams/treatment; *p<0.05, ** p<0.01, ***p>0.001 as determined by Kruskal-Wallis (KW) one-way ANOVA or Kolmogorov-Smirnov (KS) test.

| **Placenta** | | | |
| --- | --- | --- | --- |
| Cytokines and Chemokines | Statistical Analysis | | |
| Type | Test | F, H or W value | P value |
| EGF | Kruskal-Wallis | F (4,26) = 4.04 | **P=0.011*** |
| Eotaxin | ANOVA | F (4, 30) = 5.47 | **P=0.0020*** |
| Fracktalkine (CX3CL1) | ANOVA | F (4, 30) = 0.48 | P=0.74 |
| G-CSF | ANOVA | F (4, 30) = 3.32 | **P=0.023*** |
| GM-CSF | ANOVA | F (4, 29) = 0.39 | P=0.81 |
| GRO/KC (CXCL-1) | Kruskal-Wallis | H (4) =6.57 | P=0.16 |
| IFNy | Welch's | W(4.00, 13.05)=10.66 | **P=0.00050*** |
| IL-1α | Kruskal-Wallis | H (4) =20.72 | **P=0.00040*** |
| IL-1β | ANOVA | F (4, 29) = 13.08 | **P<0.00010*** |
| IL-2 | ANOVA | F (4, 30) = 1.69 | P=0.18 |
| IL-4 | Kruskal-Wallis | H (4) =12.46 | **P=0.014*** |
| IL-5 | ANOVA | F (4,30 ) = 1.88 | P=0.49 |
| IL-6 | ANOVA | F (4, 30) = 57.78 | **P<0.00020*** |
| IL-10 | ANOVA | F (4, 40) = 3.23 | **P=0.025*** |
| IL-12p70 | ANOVA | F (4, 30) = 1.66 | P=0.19 |
| IL-13 | ANOVA | F (4, 30) = 1.68 | P=0.18 |
| IL-17A | ANOVA | F (4, 30) = 14.03 | **P<0.00010*** |
| IL-18 | ANOVA | F (4, 30) = 1.49 | P=0.23 |
| IP-10 (CXCL10) | Kruskal-Wallis | H (4) = 12.3 | **P=0.015*** |
| Leptin | ANOVA | F (4, 30) = 0.51 | P=0.73 |
| LIX (CXCL5) | Welch's | W(4.00, 14.04)=5.16 | **P=0.047*** |
| MCP-1 (CCL-2) | Kruskal-Wallis | H (4) = 4.16 | P=0.39 |
| MIP-1α (CCL3) | Kruskal-Wallis | H (4) = 20.13 | **P=0.00050*** |
| MIP-2 (CXCL2) | ANOVA | F (4, 28) = 1.29 | P=0.30 |
| RANTES (CCL5) | ANOVA | F (4, 28) = 3.47 | **P=0.020*** |
| TNFα | ANOVA | F (4, 29) = 4.20 | **P=0.0083*** |
| VEGF | ANOVA | F (4, 30) = 1.73 | P=0.17 |

**Table S1.** Statistical analysis for placenta multiplex assay.

| **Combined Male and Female Fetal Brains** | | | |
| --- | --- | --- | --- |
| Cytokines and Chemokines | Statistical Analysis | | |
| Type | Test | F, H or W value | P value |
| EGF | Kruskal-Wallis | H (4) = 13.55 | **P=0.0089*** |
| Eotaxin | Welch's | W (4.000, 29.72)=1.635 | P=0.19 |
| Fracktalkine (CX3CL1) | Kruskal-Wallis | H (4) =1.59 | P=0.81 |
| G-CSF | ANOVA | F (4, 58)= 0.72 | P=0.58 |
| GM-CSF | Kruskal-Wallis | H (4) = 4.61 | P=0.33 |
| GRO/KC (CXCL-1) | Kruskal-Wallis | W (4.000, 26.85)=3.64 | P=0.021 |
| IFNy | ANOVA | F (4, 61) = 6.00 | **P=0.00040*** |
| IL-1α | ANOVA | F (4, 60) = 3.19 | **P=0.019*** |
| IL-1β | Welch's | (4.000, 27.14)=5.37 | **P=0.0026*** |
| IL-2 | ANOVA | F (4, 61) = 4.27 | **P=0.0041*** |
| IL-4 | ANOVA | F (4, 61) = 6.10 | **P=0.00030*** |
| IL-5 | ANOVA | F (4, 61) = 0.73 | P=0.57 |
| IL-6 | Welch's | W (4.00, 39.43)=9.13 | **P<0.00010*** |
| IL-10 | Kruskal-Wallis | H(4) = 21.26 | **P=0.0003*** |
| IL-12p70 | ANOVA | F (4, 61) = 0.62 | P=0.65 |
| IL-13 | ANOVA | F (4, 60) = 0.59 | P=0.67 |
| IL-17A | ANOVA | F (4, 61) = 3.71 | **P=0.0091*** |
| IL-18 | Welch's | W (4.00, 29.19)=4.84 | **P=0.0041*** |
| IP-10 (CXCL10) | Welch's | W (4.00, 40.52)=4.05 | **P=0.0075*** |
| Leptin | ANOVA | F (4, 61) = 0.57 | P=0.68 |
| LIX (CXCL5) | Welch's | W(4.00, 29.31)=3.61 | **P=0.017*** |
| MCP-1 (CCL-2) | Kruskal-Wallis | H (4) = 0.82 | P=0.93 |
| MIP-1α (CCL3) | Welch's | W (4.00, 30.24)=5.10 | **P=0.0029*** |
| MIP-2 (CXCL2) | Kruskal-Wallis | H (4) = 1.70 | P=0.79 |
| RANTES (CCL5) | Welch's | W (4.00, 29.92)=7.02 | **P=0.0015*** |
| TNFα | Welch's | W (4.00, 28.33)=6.84 | **P=0.00060*** |
| VEGF |  | F (4, 61) = 2.80 | **P=0.034*** |

**Table S2. Statistical analysis for male and female fetal brain multiplex assay.**
